# Genome-wide screen of cell-cycle regulators in normal and tumor cells identifies a differential response to nucleosome depletion

**DOI:** 10.1101/060350

**Authors:** Maria Sokolova, Mikko Turunen, Oliver Mortusewicz, Teemu Kivioja, Patrick Herr, Anna Vähärautio, Mikael Björklund, Minna Taipale, Thomas Helleday, Jussi Taipale

## Abstract

To identify cell cycle regulators that enable cancer cells to replicate DNA and divide in an unrestricted manner, we performed a parallel genome-wide RNAi screen in normal and cancer cell lines. In addition to many shared regulators, we found that tumor and normal cells are differentially sensitive to loss of the histone genes transcriptional regulator CASP8AP2. In cancer cells, loss of CASP8AP2 leads to a failure to synthesize sufficient amount of histones in the S-phase of the cell cycle, resulting in slowing of individual replication forks. Despite this, DNA replication fails to arrest, and tumor cells progress in an elongated S-phase that lasts several days, finally resulting in death of most of the affected cells. In contrast, depletion of CASP8AP2 in normal cells triggers a response that arrests viable cells in S-phase. The arrest is dependent on p53, and preceded by accumulation of markers of DNA damage, indicating that nucleosome depletion is sensed in normal cells via a DNA-damage-like response that is defective in tumor cells.

## Introduction

The cell cycle can be divided into two distinct periods, the interphase and the mitotic phase (M). During the interphase, cells are growing and duplicating their DNA, whereas in the mitotic phase, cells divide into two daughter cells. Interphase consists of three separate phases: gap1 (G1), DNA-synthesis (S) and gap2 (G2). Passing from one phase of cell cycle to the next is mainly regulated by cyclins and cyclin-dependent kinases. During the cell cycle, several checkpoints control that cells are ready to pass from one phase to the next. Interphase has two main checkpoints. One is in G1 phase, ensuring that everything is ready for DNA replication and the other is in G2 phase to verify that DNA replication is completed and that any damage to DNA is repaired. In addition, during the S phase, when the entire genome is replicated several checkpoint pathways can be activated as a response to DNA damage or stalled replication forks.

DNA replication is tightly coordinated with chromatin assembly, which depends on the recycling of parental histones and deposition of newly synthetized histones^1^. *Drosophila* and yeast *S. cerevisiae* cells can complete S phase without *de novo* histone synthesis^2, 3^. However, loss of histone expression or limiting assembly of nucleosomes to DNA by targeting chromatin assembly factors such as CAF-1, ASF1 and SLBP have been reported to induce S phase arrest in human tumor cells^4–8^. However, the mechanism of this arrest is still poorly understood.

Many regulators of the cell cycle have been identified by loss of function screens in yeast. Genome-wide RNAi screens have subsequently been used to identify both regulators that are conserved in and specific for higher organisms such as Drosophila^9^ and human^10–14^. In addition to differences between species, regulation of the cell cycle is often altered in normal and tumor cells from the same organism. This is thought to be at least in part due to mutations in cell cycle regulatory proteins such as RB1^15^ and p53^16^. However, a systematic comparative study that would identify regulators that are differentially required in normal and tumor cells has not been performed.

## Results

### Genome-wide RNAi screen identifies differential regulation of S-phase progression in normal and cancer cells

To understand similarities and differences in cell cycle regulation in normal and cancer cells, we performed a genome-wide RNAi screen simultaneously in two distinct cell lines: the osteosarcoma cell line U2OS and the hTERT-immortalized normal retinal pigment epithelial cell line hTERT-RPE1.

U2OS and hTERT-RPE1 cells grown on 96-well plates were transfected in triplicate with the same transfection mixes, containing pooled siRNAs (Qiagen) targeting a single gene per well. A total of 23 348 genes were targeted, and the cell cycle phase of the cells was analyzed after 3 days using laser scanning cytometry (**Fig. 1a**). For 17 095 targeted genes a total of five phenotypes were analyzed: the total number of cells, the fraction of cells in G1, S and G2 phases, and the fraction of cells that had higher DNA content than G2 phase cells (overG2; **Fig. 1b, Supplementary Table S1**). These represent cells that have replicated DNA more than once without dividing.

**Figure 1:**
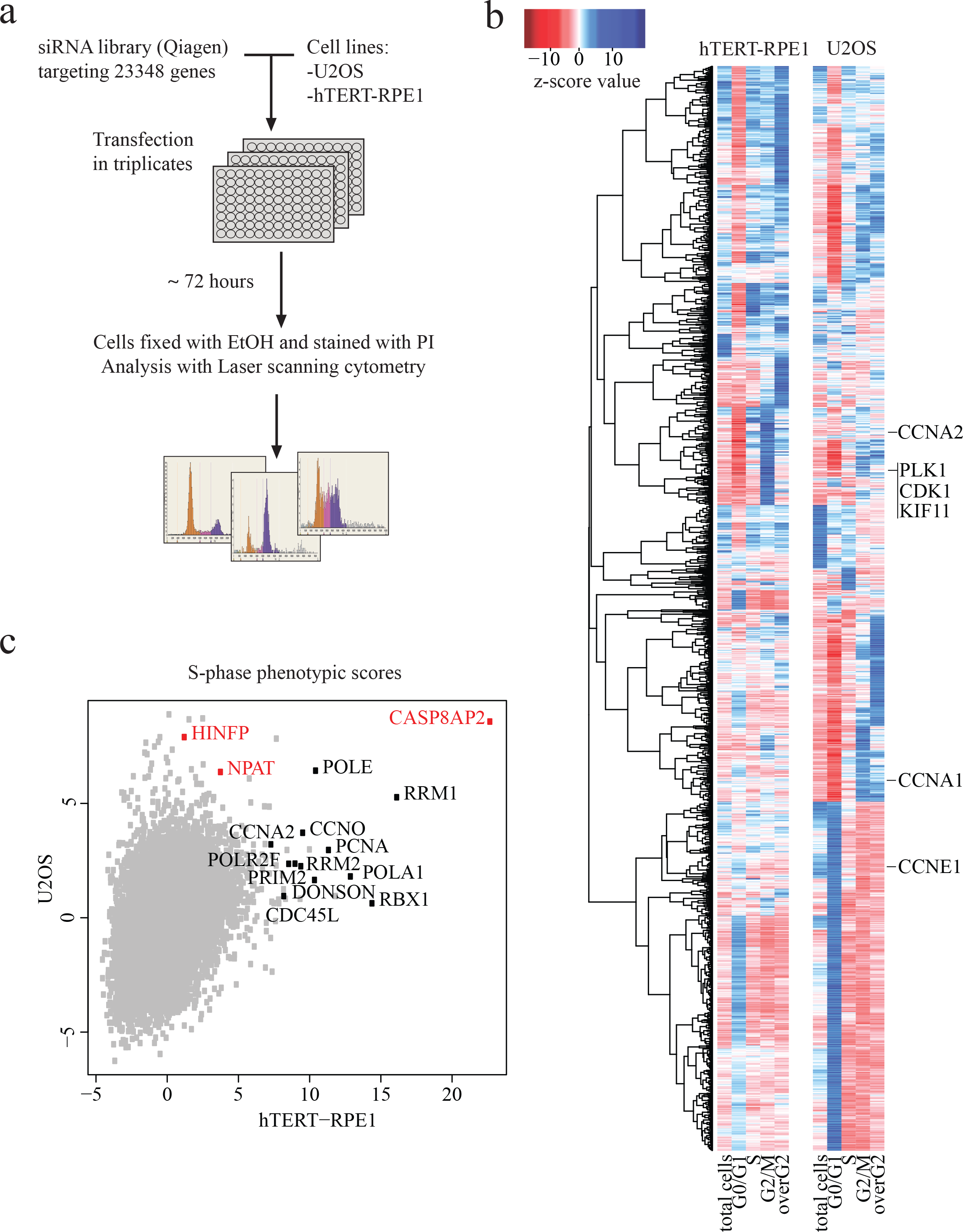
Genome-wide RNAi screen comparing normal and tumor cells. **a**, Schematic representation of genome-scale RNAi screening in human immortalized (hTERT-RPE1) and cancer (U2OS) cell lines. **b**, Hierarchical clustering of cell cycle regulators (z-score for at least one parameter is >5 or <-5) based on their phenotypic scores (for full data, see Supplementary Table S1). Note that known genes required for cell division (e.g. CDK1, KIF11, PLK1) cluster together for both cell lines. **c**, S-phase z-scores in hTERT-RPE1 and U2OS cell lines. Most S-phase arrested hits for hTERT-RPE1 are genes involved in DNA replication and S-phase progression (marked black). Transcriptional regulators of histone genes are marked in red.

In general, RNAi treatments targeting genes known to be required for mitosis (e.g. *CDK1, PLK1, KIF11*) had similar phenotypes in both cell lines (**Fig. 1b, Supplementary Fig. 1a**). However, several known S-phase genes (e.g. *PCNA, POLA1, RBX1, RRM1*) had much stronger S phase arrested phenotypes in hTERT-RPE1 cells compared to U2OS (**Fig. 1c**). One of the strongest S-phase phenotypes in both cell lines was caused by siRNAs targeting caspase 8 associated protein 2 (*CASP8AP2*, also known as *FLASH*; **Fig. 1c**). *CASP8AP2*was also the strongest S-phase regulator in a secondary screen with a Dharmacon siRNA library targeting 55 of the identified cell cycle genes in nine different cell lines (**Supplementary Table S2; Supplementary Fig. 1b**). siRNA targeting of two other known regulators of histone gene transcription, *NPAT* and *HINFP* also resulted in an increase in the fraction of cells in the S-phase in most of the nine cell lines studied.

### Loss of histone gene transcription regulators differentially affects S-phase progression

To validate disruption of S-phase progression by loss of the regulators of histone genes we transfected U2OS and hTERT-RPE1 cells with *CASP8AP2, NPAT, HINFP* and control siRNA pools and then measured the DNA synthesis rate by incorporation of the thymidine analogue 5-Ethynyl-2′-deoxyuridine (EdU). In both U2OS and hTERT-RPE1 cells, knockdown of *CASP8AP2* dramatically decreased EdU incorporation in S-phase. Knockdown of *NPAT* and *HINFP* had a similar effect in U2OS cells with accumulation of cells with poor EdU incorporation. However, in hTERT-RPE1 cells depletion of *NPAT* and *HINFP* failed to appreciably affect S-phase progression (**Fig. 2a**).

**Figure 2:**
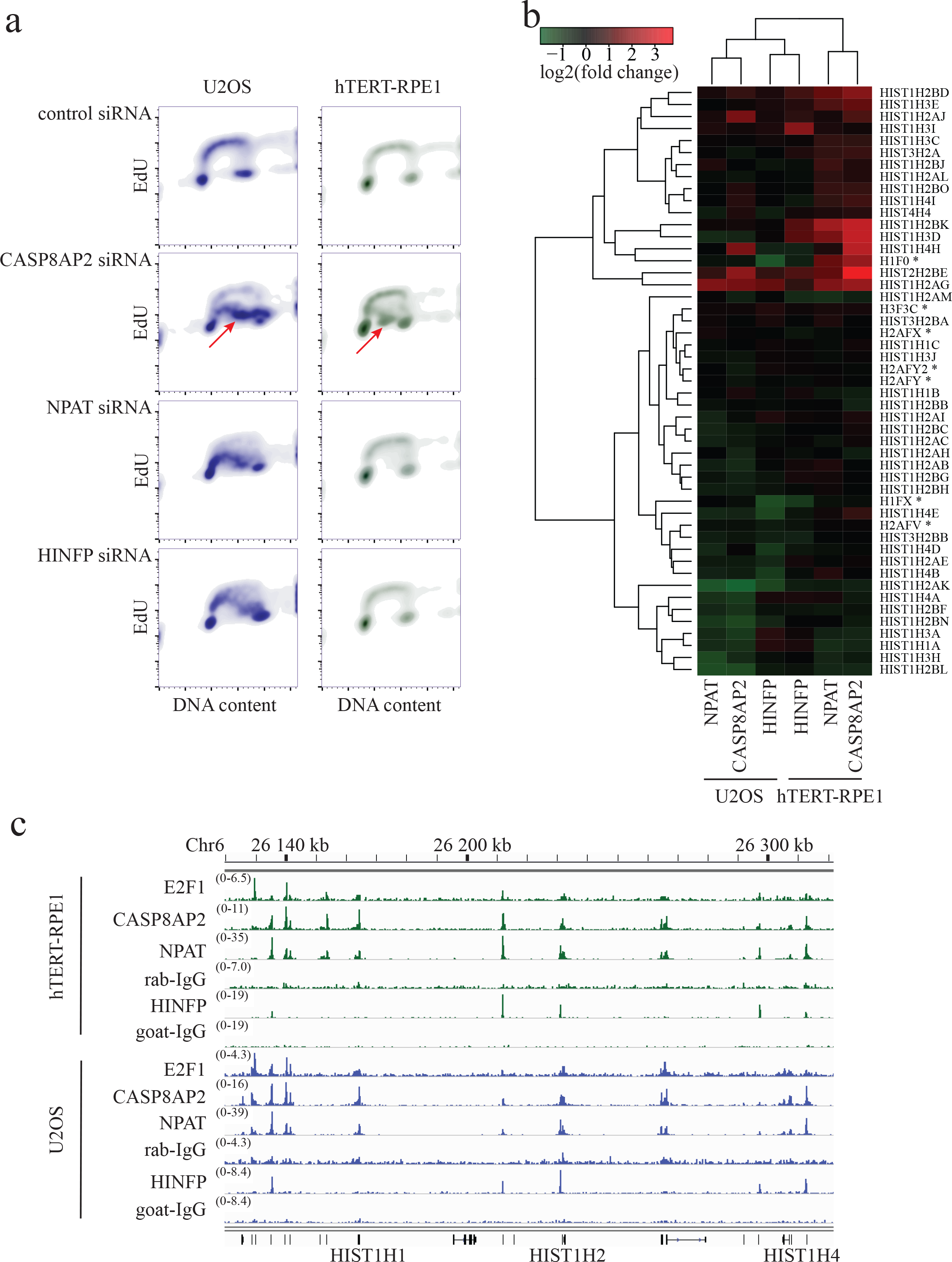
Regulation of DNA replication and expression of histone genes by CASP8AP2, NPAT and HINFP. **a**, Flow cytometric analysis of DNA content (x-axis) and DNA replication (EdU incorporation; y-axis) shows partial or complete DNA replication progression three days after knockdown of CASP8AP2, NPAT and HINFP in tumor (U2OS) and normal (hTERT-RPE1) cells. Note that in both cell lines, CASP8AP2 RNAi results in formation of a population of S-phase cells with low EdU incorporation (red arrowheads). **b**, Analysis of expression of histone genes following knockdown of the indicated genes in U2OS and hTERT-RPE1 cells. Replication-independent histone genes are marked with an asterisk. **c**, Location-analysis of transcriptional regulators at histone gene cluster on chromosome 6p22. Cell lines and antibodies used in ChIP-Seq are indicated on the left, and signal intensity as number of reads is shown in parentheses above each track. Note that CASP8AP2 and NPAT co-bind to transcription start sites of replication-dependent histone genes (bottom) in this cluster.

### CASP8AP2, NPAT, HINFP and E2F1 have different impact on histone gene expression

To determine the effect of loss of CASP8AP2, NPAT and HINFP on histone gene expression, we profiled gene-expression in siRNA treated U2OS and hTERT-RPE1 cells using Affymetrix WT1.1 arrays (**Supplementary Table S3**). We found that CASP8AP2, NPAT and HINFP do not regulate expression of each other, but mainly affect the expression of histone genes. Most histone genes were downregulated in U2OS cells following loss of CASP8AP2,
NPAT or HINFP. In normal cells, some highly expressed histone genes were downregulated, albeit less than in tumor cells. In addition, many histone genes that are normally expressed at lower levels were upregulated (**Fig. 2b; Supplementary Table S3**).

To identify whether CASP8AP2, NPAT and HINFP directly bind to the histone gene promoter regions we carried out ChIP-Seq in U2OS and hTERT-RPE1 cells. Consistent with previous findings, HINFP was found enriched near transcription start sites (TSSs) of replication-dependent histones H4 and H2B^17–20^ (**Supplementary Table S4 and S5**). We also found that HINFP regulated two replication-independent histone H1 genes, H1F0 and H1FX (**Supplementary Table S4 and S5**). In contrast, CASP8AP2 and NPAT ChIP-Seq peaks were only found colocalized at replication-dependent histone genes on chromosomes 1, 6 and 12 in both cell lines (**Fig. 2c**, **Supplementary Table S4 and S5**). These results indicate that CASP8AP2 and NPAT regulate only replication-dependent histones, whereas HINFP regulates a subset of replication dependent histones (H4 and H2B), and two replication indipendent H1 variants (H1F0 and H1FX).

Another histone gene regulator, E2F1^21, 22^, also bound to TSSs of many histone genes, including both replication dependent and independent histones (**Supplementary Table S4 and S5**). In addition, E2F1 bound to the promoter of CASP8AP2, suggesting that E2F proteins control CASP8AP2 and histone expression directly and via a feed-forward loop, respectively.

### CASP8AP2 knockdown results in low histone H3 protein levels and slows progression of replication forks in osteosarcoma cells

To analyze the long-term effect of CASP8AP2 loss on S-phase progression and histone protein levels, we treated U2OS and hTERT-RPE1 cells with CASP8AP2 siRNAs, and analyzed DNA content, histone H3 protein level, and EdU incorporation by flow cytometry in the same population of the cells. We found that CASP8AP2 siRNA treatment did not completely arrest U2OS cells in S-phase, but instead dramatically slowed down S-phase progression, resulting in an S-phase that lasted more than 3 days (**Fig. 3a**). The slowdown in S-phase was accompanied by increased cell death, and a marked decrease in histone H3 protein levels in the surviving cells that continued to replicate (**Fig. 3b**). In contrast, viable hTERT-RPE1 cells were arrested in S-phase by CASP8AP2 siRNA, and very little if any effect on histone H3 protein levels was detected (**Supplementary Fig 2a-d**).

**Figure 3:**
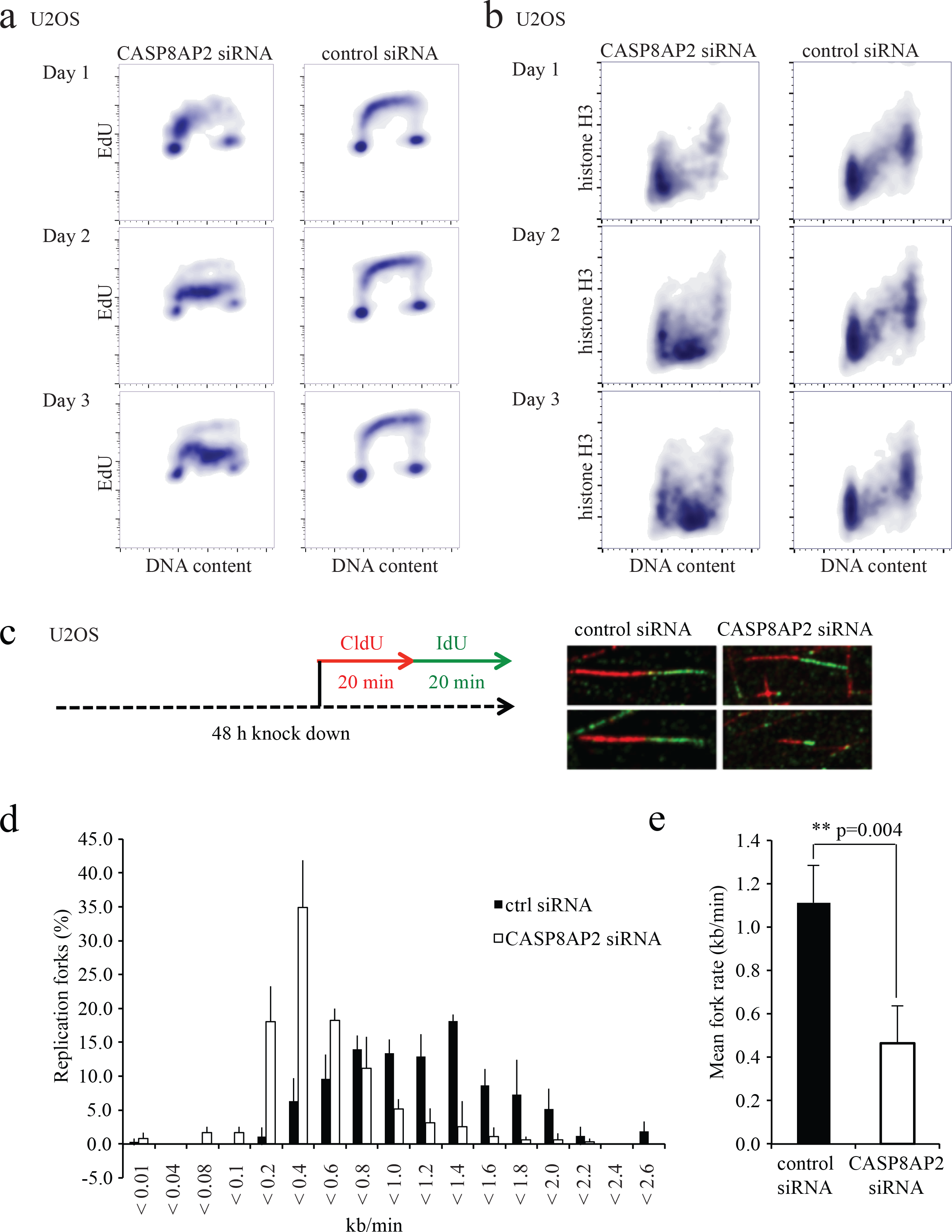
Tumor cells continue to progress in S-phase despite low nucleosome levels induced by CASP8AP2 knockdown. **a, b**, Flow cytometry analysis of DNA content, DNA replication and histone H3 levels in siRNA treated cells. Note that tumor cells continue to replicate their DNA slowly (a) for multiple days, despite low amount of histone H3 (b). **c**, Schematic representation of the DNA fiber assay measuring the length of labeled DNA strands. Images show representative DNA fibers 2d after control or CASP8AP2 siRNA transfection. **d**, Distribution of replication fork speeds in control and CASP8AP2 depleted cells. **e**, Significant reduction of mean fork rate in CASP8AP2 siRNA treated cells. Mean of three independent experiments are shown. Error bars represent one standard deviation.

To identify the mechanism of the slowdown of the cell cycle in the U2OS cells, we analyzed the speed of replication forks in CASP8AP2 siRNA treated U2OS cells using a DNA fiber assay, where DNA is labelled by two different labels consecutively, and then analyzed visually to detect the length of the labelled regions. This analysis confirmed earlier observations^23^ that CASP8AP2 siRNA generally decreases the speed by which replication forks progress (**Fig. 3c–e**), suggesting that in human cancer cells, individual replication forks are affected by nucleosomes loading behind them.

### The ability of normal cells to activate H2AX phosphorylation in response to deregulation of histone expression is p53 dependent

To determine why normal and tumor cells respond differentially to CASP8AP2 loss, we examined the expression profiles of non-histone genes after CASP8AP2, NPAT and HINFP siRNA treatment. This analysis revealed that p53 target genes were upregulated in hTERT-RPE1, but not in p53-proficient U2OS cells (**Fig. 4a**). The p53 target genes were upregulated relatively late, clearly, after the changes in histone gene expression were observed (**Supplementary Fig. 3 a,b**; **Supplementary Table S6**).

**Figure 4:**
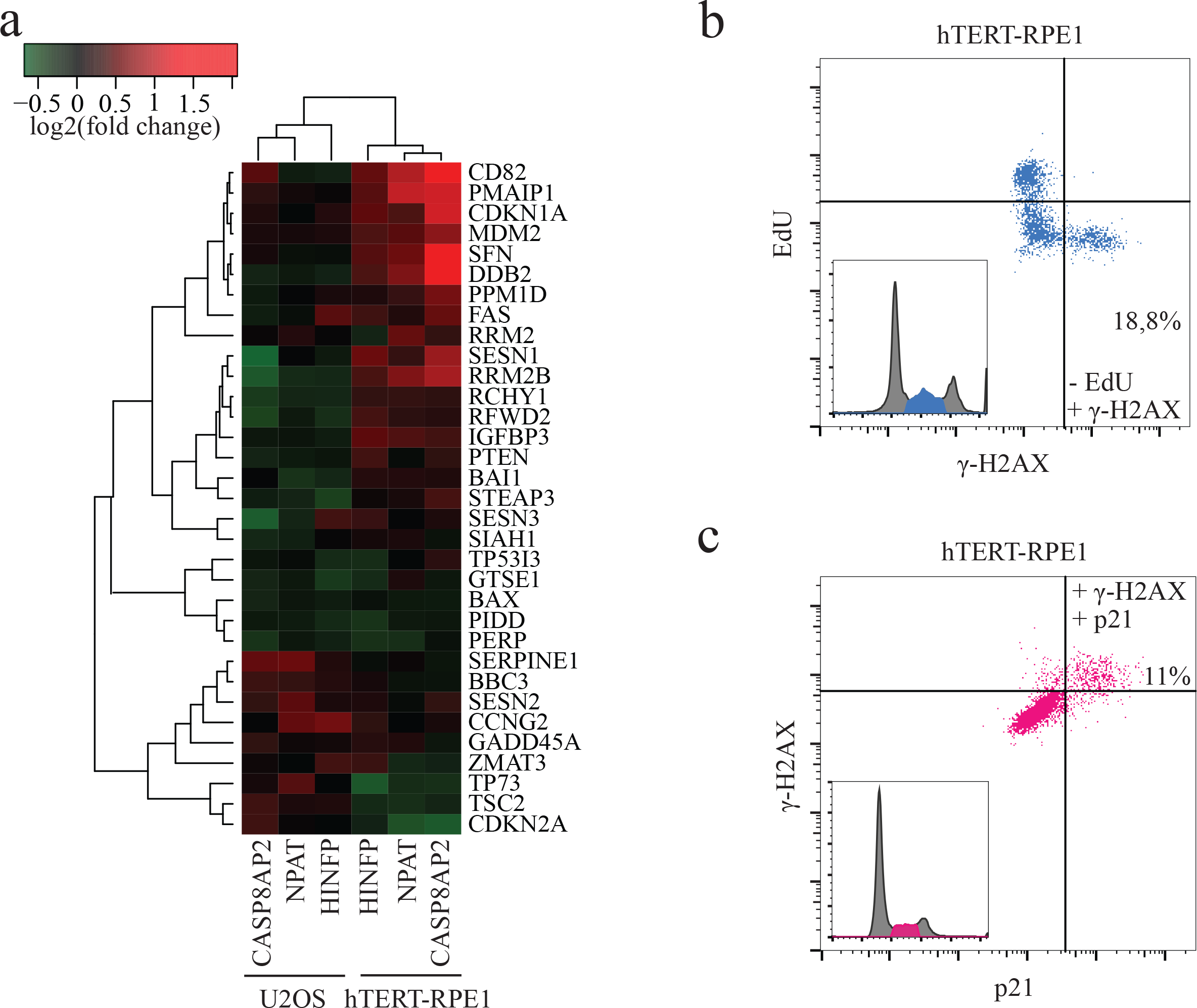
CASP8AP2 siRNA triggers activation of a p53 dependent S-phase checkpoint in normal but not in tumor cells. **a**, Analysis of p53 target gene expression three days after knockdown of the transcriptional regulators of histone genes indicated. **b**, Flow cytometry analysis of CASP8AP2 siRNA treated hTERT-RPE1 cells indicates that non-replicated cells arrested in S-phase express a marker for DNA damage signaling (γ-H2AX; low left hand corner). **c**, Cells with activated DNA damage signaling have activated p53, based on expression of the p53 target-gene p21(upper left hand corner).

One possibility that would explain the activation of p53 is that the slow replication causes DNA damage or a DNA damage-like state that is sensed by p53. Consistently with this hypothesis, an increase in a marker commonly associated with DNA damage, gamma-H2AX, was observed more in hTERT-RPE1 cell line (**Fig. 4b; Supplementary Fig. 3c,d**) and less in U2OS cell line (**Supplementary Fig. 3c,d; 4a-d**). To address whether the p53-dependent pathway is responsible for the S-phase arrest observed in normal but not in tumor cells, we analyzed the expression of the p53 target gene p21. We found that after CASP8AP2 RNAi treatment, p21 protein was induced more in hTERT-RPE1 cells (**Supplementary Fig. 5a**), and only in S-phase cells that express elevated gamma-H2AX (**Fig. 4c**). Similar results were found in two other normal human cell lines, HFL1 and CCD-1112Sk (**Supplementary Fig. 6a-d**).

To test if the effect is dependent on p53, we generated a derivative of hTERT-RPE1 cell line that lacks p53 using CRISPR/Cas9 genome engineering. This cell line failed to completely arrest in S-phase or induce p21 in response to CASP8AP2 RNAi (**Supplementary Fig. 5a,b**). In addition, significantly lower levels of gamma-H2AX accumulation were observed in the p53 deficient cell line (**Supplementary Figs. 3c,d and 6a-d**). The response of the p53 deficient cell line was similar to that of tumor cells, including U2OS and HCT116 (**Supplementary Fig. 6a-d**).

## Discussion

Genome-wide RNAi screens have become very powerful and informative approaches for the analysis of cell cycle regulation and for identification of new regulators of cell cycle progression and checkpoint control. Unlikely previously published genome-wide RNAi screens performed to identify genes affecting cell cycle and cell size in *Drosophila* cells^9^ and in human cancer cells^10–14^ the present study compares two distinct cell lines: the osteosarcoma cell line U2OS and the hTERT-immortalized normal retinal pigment epithelial cell line hTERT-RPE1. As expected, several known mitotic regulators, and the known transcriptional regulator of histones, CASP8AP2 displayed strong phenotypes in both types of cells. However, some DNA-replication regulators, including two other transcriptional regulators of histones, HINFP and NPAT, displayed differential effects in the two cell types.

The canonical histones are exclusively expressed in S phase of cell cycle^22, 24, 25^. This process is regulated by E2F and cyclin E-Cdk2 kinase through phosphorylation of NPAT^19, 21, 26-28^. Based on microarray data and flow cytometry, the depletion of CASP8AP2 caused strong downregulation of replication-dependent histones in U2OS cells. In normal cells, loss of CASP8AP2 also resulted in deregulation of histone expression, resulting in both positive and negative effects at the level of individual histone genes. The transcriptional regulation of histone genes by CASP8AP2 and NPAT is likely direct, based on their known interaction^29^, and our chromatin immunoprecipitation followed by sequencing (ChIP-seq) results showing localization of NPAT and CASP8AP2 almost exclusively to the replication-dependent histone loci.

In order to understand the mechanism behind the differential response of normal and tumor cells, we analyzed the S-phase phenotypes in more detail, using EdU incorporation assays in multiple tumor and normal cells depleted of CASP8AP2. This analysis revealed that the increase in cells in the S-phase observed in tumor cells after CASP8AP2 loss was not caused by an S-phase arrest. Instead, it resulted from a dramatic slowdown of DNA replication. In contrast, the S-phase progression of normal cells was arrested. Further analysis revealed also that the deregulation of nucleosome gene expression triggers a p53-dependent pathway in normal cells, but not in U2OS tumor cells. Thus, p53 activation is a consequence, not the cause, of the observed DNA replication phenotype.

One possibility that would explain the activation of p53 is that imbalance of histone proteins and the disruption of replication process causes DNA damage or a DNA damage-like state that is sensed by p53. Consistently with this hypothesis, an increase in a marker commonly associated with DNA damage, gamma-H2AX, was observed in hTERT-RPE1 cells following CASP8AP2 knockdown. Much lower levels of the marker were seen in similarly treated U2OS cells. Thus, downregulation of histones in tested cancer cells could cause slow progression through S-phase without sufficient accumulation of DNA damage marker to activate p53-dependent cell cycle arrest. It consistent with previous result where short-term depletion of histone genes by CASP8AP2 knockdown did not induce DNA damage markers^23^.

In summary, we have through a genome-wide analysis of normal and tumor cells identified a significant difference in regulation of cell cycle arrest in response to abnormal regulation of histone genes transcription. In both normal and tumor cells, the depletion of CASP8AP2 alters mainly the expression of replication-dependent histones. Whereas hTERT-RPE1 cells respond by triggering a p53-dependent checkpoint, which leads to cell cycle arrest and recovery, U2OS cells continue to progress in S-phase, leading to cell death. The difference was observed in all tested cell types, including eight tumor cell lines and three different normal cell types. Further studies are necessary for determining the precise mechanisms leading to cell death in cells that continue to progress in S-phase despite lack of sufficient levels of histones. The identified defect in sensing histone expression levels could also in part explain the abnormal nuclear morphology commonly associated with cancer cells. Furthermore, the identified novel vulnerability of cancer cells can potentially be used for therapeutic purposes in the future.

## SUPPLEMENTARY FIGURE LEGENDS

**Supplementary Figure 1: Genome-wide RNAi screen and secondary RNAi screen in several cancer cell lines**. **a**, Genome-wide RNAi screen identified many common regulators that decrease G1 content in human immortalized (hTERT-RPE1) and cancer (U2OS) cell lines. Samples (black circles) and three groups of controls (colored circles) are shown. Known cell cycle regulators are indicated in black typeface. **b**, Secondary RNAi screen in different cancer cell lines shows significant S-phase arrest following CASP8AP2 knockdown in all cell lines; NPAT and HINFP knockdown have similar albeit weaker effect in most of cell lines.

**Supplementary Figure 2: S-phase slow progression in hTERT-RPE1**. **a, b**, Flow cytometry analysis of DNA content, DNA replication (EdU staining) and histone H3 levels in siRNA treated hTERT-RPE1 cells. Note that normal cells arrest in S-phase (a), and do not show strong decrease in histone H3 levels (b). **c**, Western-blot analysis of H3 following subcellular protein purification indicates that most of the H3 in both cells lines is chromatin-bound. **d**, Western blot analysis confirms decreased level of histone H3 after CASP8AP2 knockdown in U2OS but not in hTERT-RPE1 cell line.

**Supplementary Figure 3: H2AX Ser139 phosphorylation and p53 independent regulation of histone gene expression and DNA replication following CASP8AP2 knockdown. a, b**, Histone gene deregulation precedes activation of p53. Microarray data in hTERT-RPE1 cells during three days following CASP8AP2 knockdown shows deregulation of (a) histone gene expression from the first day and (b) most of p53 target genes only after second day. **c**, H2AX Ser139 phosphorylation in S-phase arrested cells following CASP8AP2 knockdown in U2OSand hTERT-RPE1 cell lines with different p53 status 3 d after transfection. Note that DNA damage signaling is strongest in hTERT-RPE1 p53 wt cells. **d**, Different accumulation of γ-H2AX in CASP8AP2 depleted cancer and normal cells with different p53 status. Mean and standard deviation for biological triplicates are shown. Note that DNA damage signaling is highest in normal p53 wt cells at 3 days, decreasing at later time points. In contrast, in tumor cells and normal cells lacking p53 the response is initially weaker, but continues to increase at least up to 5 days.

**Supplementary Figure 4: CASP8AP2 is required to maintain genome integrity in U2OS**. (**a**,**a**’) The dramatic defect in S-phase progression in CASP8AP2 deficient cells is comparably strong to hydroxyurea treatment as determined by EdU incorporation. (**b**, **b**’) Loss of CASP8AP2 results in accumulation of ssDNA (RPA foci), and increased DNA damage signaling measured by the recruitment of γ-H2AX (**c**, **c**’) and p53BP1 (**d**, **d**’). For all experiments (n=3) means and s.e.m are plotted and representative images are shown. P-values (*) are calculated with Student’s t-test and scale bars represent 50 μm.

**Supplementary Figure 5: p21 activation and DNA replication progression in CASP8AP2 depleted cells**. **a**, Normal cells with wild-type p53 respond to CASP8AP2 knockdown by upregulating p21. Panels show flow cytometry analysis of p21 protein expression following CASP8AP2 knockdown in U2OS and hTERT-RPE1 cell lines with different p53 status. **b**, hTERT-RPE1 cells lacking p53 continue to replicate their DNA despite knockdown of CASP8AP2. EdU cell proliferation assay indicates that more DNA synthesis occurs in late S-phase hTERT-RPE1 p53 KO than hTERT-RPE1 p53 wt cells. Percentage of the cells with low EdU incorporation in mid-S and late-S phases is indicated black and red, respectively.

**Supplementary Figure 6: Correlation of DNA replication progression, p21 accumulation and H2AX Ser139 phosphorylation in CASP8AP2 depleted cells. a**, Number of cells in S-phase, EdU negative cells in S-phase, γ-H2AX cells in S-phase and p21 positive cells following CASP8AP2 knockdown in seven different cell lines. Mean and standard deviation for biological triplicates are shown. **b**, **c**, **d**, The correlation of amount of cells, EdU negative cells, γ-H2AX and p21 in S-phase for seven different cell lines (normal cells with p53wt are shown in brown). **b**, Accumulation of γ-H2AX in S-phase is higher for normal cell lines with p53wt than in cancer cells or normal cells with p53KO. **c**, Accumulation of γ-H2AX is higher in non-replicating or slowly replicating cells for normal cell lines with p53wt as compared to cancer cells or normal cells lacking p53. **d**, Only normal cells with wild-type p53 strongly upregulate p21 expression in S-phase in response to CASP8AP2 knockdown.

### Methods

#### Cell culture

U2OS human osteosarcoma (from M.Laiho lab, University Helsinki), HeLa human cervix adenocarcinoma (from M.Laiho lab, University Helsinki), HT1080 human fibrosarcoma (from ATCC, CRL-121), PC3 prostate adenocarcinoma (from Gonghong Wei, University Oulu), SK-N-MC human brain neuroepithelioma (from Gonghong Wei, University Oulu), SW480 colorectal adenocarcinoma (from Gonghong Wei, University Oulu) cells were cultured in DMEM supplemented with 10% fetal bovine serum and antibiotics (100 units/ml penicillin and 100 μg/ml streptomycin). The hTERT-immortalized retinal pigment epithelial hTERT-RPE1 (from ATCC, CRL-4000) and hTERT-RPE1 p53KO cells lines were cultured in F12:DMEM(1:1) supplemented with 10% fetal bovine serum and antibiotics (100 units/ml penicillin, 100 μg/ml streptomycin and 10 μg/ml hygromycin B). SAOS-2 human osteosarcoma (from M.Laiho lab, University Helsinki) were cultured in McCoy’s 5A medium supplemented with 10% fetal bovine serum and antibiotics (100 units/ml penicillin and 100 |ig/ml streptomycin). SJCRH30 human rhabdomyosarcoma cells (from ATCC, CRL-2061) were cultured in RPMI-1640 supplemented with 10% fetal bovine serum and antibiotics (100 units/ml penicillin and 100 μg/ml streptomycin).

#### Chromatin immunoprecipitation followed by sequencing (ChIP-seq)

ChIP assays were performed as previously described.^30^ Immunoprecipitations were carried out with 5 μg antibody against CASP8AP2 (sc-9088, Santa Cruz Biotechnology), NPAT (sc-67007, Santa Cruz Biotechnology), HINFP (sc-49818, Santa Cruz Biotechnology), E2F1 (sc-193, Santa Cruz Biotechnology), normal rabbit IgG (sc-2027, Santa Cruz Biotechnology) and normal goat IgG (sc-2028, Santa Cruz Biotechnology).

ChIP libraries were prepared for Illumina Genome Analyzer or HiSeq2000 sequencing as described before. ^30^ Sequencing reads were mapped to the human genome (NCBI36) and peaks called as described.^31^ Lists of all significant peaks (p > 0.00001, fold change > 2) are in Supplementary Table S4.

#### Laser scanning cytometry: Genome-wide RNAi screen

A Qiagen Human Whole Genome siRNA Set V4.0 was used for transfection in U2OS and hTERT-RPE1 cell lines. Qiagen siRNA library contains four pooled siRNAs in total amount of 0. 5 nmol siRNA per well targeting 23348 different genes (EntrezGene ID). The complete set contains a total of 308 X 96-well plates with each plate containing 80 wells with siRNA constructs as well as following controls: Qiagen negative control siRNA, siRNA against GFP (Qiagen), 3 wells with transfection reagent and 3 wells with siRNA against CDK1 (custom made positive control purchased from MWG).

For reverse transfection 4000 cells were plated in the growth medium into 96-well plates having 15 nM siRNA and 0,7 μl HiPerFect (Qiagen) in total volume of 125 μl. Transfection was performed in triplicates simultaneously for U2OS and hTERT-RPE1 cells in 16 batches. After 70-72 hours incubation cells were washed with PBS, fixed with 70% ice cold ethanol o/n and stained with PBS containing 30 μg/ml PI (Sigma-Aldrich) and 30 μg/ml RNase A (MACHEREY-NAGEL). The stained cells were scanned with an Acumen eX3 microplate cytometer (TTP LabTech).

The number of the cells in each sub-population of the cell cycle was calculated with Acumen eX3 software. Single cell population was specified based on area, depth and width parameters of the signal for each object. The DNA content histograms were gated manually for each batch to specify sub-G1-, G0/G1-, S-, G2/M-and over-G2-phase populations. Total cell number and percentages of the cells in each population were normalized two times: first using median of a plate and then median of the well position in all plates (positive controls were excluded from the normalization); then Z-score were calculated for each parameter from the median of triplicates and median of all negative controls (i.e. negative control siRNA and GFP siRNA): Z=(x-μ)/ σ, where x is the median of triplicates for each parameter; n is the median of all negative controls; σ is the standard deviation of all negative controls.

For the final data analysis the siRNA sequences were mapped to human transcripts (Ensembl version 52, genome assembly NCBI36) using bowtie version 0.11.3. In total 79534 (81.36%) siRNAs were mapped to at least one transcript without mismatches and 17095 (73%) wells have 4 siRNAs mapping the same single target gene. After normalization only data from the 17095 samples were used for further analysis. To identify hits +/–5 z-score cutoff was applied for all calculated parameters (sub-G1, G0/G1, S, G2/M and over-G2), later on subG1 was excluded from the analysis as non-informative.

Data analysis and visualization was performed in R using hopach (R package version 2.20.0; http://www.stat.berkeley.edu/~laan/, http://docpollard.org/) ^32^, gplots packages and GOstat (http://gostat.wehi.edu.au/)^33^.

### Secondary screen

A Dharmacon siGENOME SMARTpool of 91 siRNAs (including negative controls) was used for transfection in 9 different cells lines (hTERT-RPE1, U2OS, SAOS-2, SJCRH30, HeLA, PC3, SK-N-MC, SW480 and HT1080). Transfection, screening and data analysis was carried out in the same way as with the Qiagen library screen.

### Microarray data

Sets of four FlexiTube GeneSolution siRNAs (Qiagen) targeting CASP8AP2, NPAT, HINFP or AllStars Negative Control (Qiagen) were used. For reverse transfection protocol, cells at 30% confluence were plated in the growth medium into 6-well plates using 15 nM siRNA and 7 μl HiPerFect (Qiagen) in final total volume 2 100 μl. Transfection was performed in duplicates simultaneously for each siRNA for U2OS and hTERT-RPE1 cell lines. After 70-72 hours growing cells were collected and RNA extraction was performed with RNeasy kit (Qiagen) including DNase treatment according to manufacturer’s protocol. RNA was analyzed using Affymetrix Human WT 1. 1 arrays in the core facility for Bioinformatics and Expression Analysis (Karolinska Institute, Stockholm, Sweden). Raw CEL data were analyzed using the Bioconductor R packages affy^34^ and limma^35^ and for the annotation custom CDF files (ENSG)^36^ was used. p53 pathway data was retrieved from KEGG (http://www.genome.jp/kegg/).

### Flow cytometry

Sets of four FlexiTube GeneSolution siRNAs (Qiagen) against CASP8AP2, NPAT, HINFP or AllStars Negative Control (Qiagen) were used for reverse transfection protocol as in microarray analysis.

Cells were incubated with 10 μM EdU (Life Technologies) for last 1 hour before harvesting and fixing with 70% ice cold ethanol on 1, 2 and 3 days after transfection. The cells were washed with 1% BSA in PBS and prepared for flow cytometry with Click-iT^®^ EdU Alexa Fluor^®^ 647 Flow Cytometry Assay Kit (Life Technologies) according to the manufacturer’s protocol.

For the flow cytometry analysis of total histone H3 cells were harvested and fixed with 70% ice cold ethanol on 1, 2 and 3 days after transfection and left overnight in −20°C. On the next day the cells were washed with 1% BSA in PBS several times and blocked in 1% BSA in PBS for at least an hour followed by overnight incubation at +4°C with histone H3 antibody (1:500, #9715, Cell Signaling Technology, or 1:500, <u>ab1791, Abcam</u>), γ-H2AX(1:500, #9718, Cell Signaling Technology), p21 (1:100, sc-6246, Santa Cruz Biotechnology) or as a control normal rabbit IgG (1:200, sc-3888, Santa Cruz Biotechnology), normal mouse IgG (1:200, sc-2025, Santa Cruz Biotechnology). The cells were washed several times with 1% BSA in PBS and incubated with secondary antibodies (anti-rabbit Alexa488 or anti-mouse Alexa488, Life Technologies) for an hour. After several washes with 1% BSA in PBS cells were stained for DNA content with 5μM DRAQ5 (BioStatus Limited) and 10 mg/ml RNaseA (MACHEREY-NAGEL) for 30 min. Flow cytometry was performed using MACSQuant Analyzer (Miltenyi Biotec) and data analyzed using FlowJo (FlowJo, LLC).

### Western Blotting Analysis

Different subcellular protein fractions from hTERT-RPE1 and U2OS cells were extracted using Subcellular Protein Fractionation Kit for Cultured Cells (78840, Pierce Biotechnology) according to the manufacturer’s protocol. Samples were prepared in Laemmli sample buffer and analyzed with 10%-SDS PAGE followed by immunoblotting analysis using the following antibodies: histone H3 (#9715, Cell Signaling Technology) and α-tubulin (T9026, Sigma-Aldrich).

For the total histone H3 level analysis sets of four FlexiTube GeneSolution siRNAs (Qiagen) against CASP8AP2 or AllStars Negative Control (Qiagen) were used for reverse transfection protocol in 6 well-plates as described in microarray analysis. 10 μg of total protein was analyzed by 10% SDS-PAGE followed by immunoblotting analysis using the following antibodies: histone H3 (#9715, Cell Signaling Technology) and α-tubulin (T9026, Sigma-Aldrich).

Detection of the antibodies was carried out with Pierce ECL plus western blotting substrate (#32132, ThermoScientific) and processed with Gel Doc™ XR+ and ChemiDoc™ XRS+ Systems with Image Lab™ 5.0 Software (Bio-Rad).

### Genome editing

Generating hTERT-RPE1 p53KO cell line using CRISPR (clustered regularly interspaced short palindromic repeats) in complex with Cas9 (CRISPR associated protein 9) protein technology was carried out according previously published protocol^37^.
The following two 23 bp guide RNAs (gRNAs) in the p53 coding region were designed using the CRISPR web tool (http://crispr.mit.edu/):

> TP53_1 in exon 4: GCATTGTTCAATATCGTCCGGGG
>
> TP53_2 in exon 2: GCGACGCTAGGATCTGACTGCGG

Selected target sequences were incorporated into two 60-mer oligonucleotides corresponding the exon 4 and exon 2 targets.

> TP53_1F for exon 4:
>
> TTTCTTGGCTTTATATATCTTGTGGAAAGGACGAAACACCGCATTGTTCAATATCGTCCG
>
> TP53_1R for exon 4:
>
> GACTAGCCTTATTTTAACTTGCTATTTCTAGCTCTAAAACCGGACGATATTGAACAATGC
>
> TP53_2F for exon 2:
>
> TTTCTTGGCTTTATATATCTTGTGGAAAGGACGAAACACCGCGACGCTAGGATCTGACTG
>
> TP53_2R for exon 2:
>
> GACTAGCCTTATTTTAACTTGCTATTTCTAGCTCTAAAACCAGTCAGATCCTAGCGTCGC

The two oligos were annealed and Supplementary for 30 min at 72°C to make a 100 bp double stranded DNA fragment using Phusion polymerase (NEB). Using Gibson assembly (NEB) reaction according to the manufacturer’s protocol the 100 bp DNA fragments were incorporated into the gRNA cloning vector (http://www.addgene.org/41824/) and the reaction products were transformed into chemically competent DH5alpha cells. Plasmid DNA from five single colonies for both constructs (TP53_1 and TP53_2) were extracted using Plasmid midiprep (Qiagen).

After sequence verification plasmids containing insertions with TP53_1 and TP53_2 gDNAs were co-transfected with hCas9 plasmid (http://www.addgene.org/41815/) into the hTERT-RPE1 cells using FuGENE^®^ HD Transfection Reagent (Promega) according to the manufacturer’s protocol.

To confirm genome editing two days after transfection cells were split and genotyped using Phusion polymerase (NEB) with following primers:

gDNA_p53_F: ACACTGACAGGAAGCCAAAGGG

gDNA_p53_R: ATCCCCACTTTTCCTCTTGCAGC

The cells with p53 KO phenotype were split in low density into several 96 well plates to obtain single-cell colonies. After several single-cell colonies had reached confluence they were split and p53 levels were verified with Western Blotting using p53 antibody (1:200, DO-1, sc-126, Santa Cruz Biotechnology) as described above. In three hTERT-RPE1 clones p53 protein was not detected and p53 status was verified with Sanger sequencing of genomic and cDNA with gDNA_p53 primers and cDNA primers:

cDNA_p53_F: CAGCCAGACTGCCTTCCG

cDNA_p53_R: GACAGGCACAAACACGCACC

Of the obtained clones, we chose to use a clone with a homozygous 1 bp insertion in exon 4 of p53, causing a frameshift mutation at codon position 48, resulting in mutation of Asp 48 to Glu, and truncation of the protein after two amino-acids out-of-frame peptide (Glu-Arg-Tyr-**Stop**).

### DNA fiber assay

2 × 10^5^ U2OS cells were reverse transfected with either 15 nM control or CASP8AP2 siRNA according to the manufacturer’s protocol (HiPerFect, Qiagen). After 48 h, cells were pulse labelled with 25 mM CldU followed by 250 mM IdU for 20 min each and harvested immediately. DNA fiber spreads were prepared as previously described^29^. In short, DNA was denatured using 2.5M HCl and incubated with rat anti-BrdU monoclonal antibody (AbD Serotec, 1:1000) and mouse monoclonal anti-BrdU antibody (Becton Dickinson, 1:750) overnight at 4°C. Objective slides were fixed in 4% formaldehyde and subsequently incubated with an AlexaFluor 555-conjugated goat anti-rat IgG and AlexaFluor 488-conjugated goat anti-mouse IgG (Molecular Probes, 1:500) for 1.5 h at room temperature. Stained DNA fibers were imaged with a Zeiss LSM710 confocal laser scanning microscope, equipped with a Plan-Apochromat 63×/1.40 Oil DIC M27 objective. Alexa 488 and 555 were excited with a 488 nm Ar laser line and a 561 nm DPSS laser line, respectively. Confocal images were recorded with a frame size of 1024×1024 pixels and a pixel size of 130 nm. CldU and IdU track length were measured using ImageJ (http://rsb.info.nih.gov/ij/) and mm values were converted into kb using the conversion factor 1mm = 2.59 kb.^38^

## Supplementary Information

Supplementary Tables S1-S6.

Supplementary Figures 1-6

## Acknowledgments

We thank Sini Miettinen for technical assistance, Drs. M. Laiho and G. Wei for for supplying several cell lines and Dr. Bernhard Schmierer for critical review of the manuscript. This work was supported by the Academy of Finland Center of Excellence in Cancer Genetics and Finnish Cancer Organizations.

## Author contributions

M.S. performed most of the experiments with help from M.Tur.
O.M. performed DNA fiber assay, P.H. performed immunofluorescence experiments. T.K. performed siRNA sequences mapping to human transcripts. M.S. carried out data analyses and interpreted the results with input from M.B., M.T., T.K. and A.V. T. H. and J.T. supervised experiments and data analysis. M.S and J.T. wrote the manuscript. M.S., M.Tur., T.K., A.V., M.B., M.T., T.H. and J.T. discussed the results and commented on the manuscript.

